# Oscillatory traveling waves during visual entrainment in autistic and neurotypical adults

**DOI:** 10.64898/2026.01.29.702587

**Authors:** Andrea Alamia, Jakob C. B. Schwenk, Johan Wagemans, Laurie-Anne Sapey-Triomphe

**Affiliations:** Cerco, CNRS, Université de Toulouse, Toulouse 31052, France; Department of Brain and Cognition, Leuven Brain Institute, KU Leuven, 3000 Leuven, Belgium; Université Claude Bernard Lyon 1, CNRS, INSERM, Centre de Recherche en Neurosciences de Lyon CRNL U1028 UMR5292, CAP team, F-69500, Bron, France

**Keywords:** traveling waves, visual entrainment, autism spectrum disorder, electroencephalography, predictive coding

## Abstract

Neural oscillations play a crucial role in cognition, and several studies emphasize the importance of considering them as traveling waves that propagate through brain regions. In scalp EEG, waves typically travel along the anterior–posterior axis: forward waves (occipital-to-frontal) predominate during sensory stimulation, while backward waves (frontal-to-occipital) emerge during rest and top-down modulation. Within the predictive coding framework, backward waves are proposed to convey predictive signals, whereas forward waves reflect sensory processes and feedforward propagation of prediction errors. In this study, we investigated traveling wave dynamics during a visual entrainment task in neurotypical (N=25) and autistic (Autism Spectrum Disorder - ASD) adults (N=24). Our results replicated previous findings in neurotypical participants, as we observed an increase in backward waves during rhythmic visual stimulation, consistent with enhanced top-down predictions. Notably, we observed the opposite pattern in the ASD group, characterized by a pronounced increase in forward waves at the entrained frequency during visual stimulation. These results align with predictive coding accounts of autistic perception, which hypothesize an imbalance between predictions and sensory evidence. Specifically, an increase in forward wave may reflect a bias toward bottom-up sensory signaling over predictive feedback, due to atypical hierarchical communication across brain regions in ASD. Together, our findings shed new light on the oscillatory dynamics involved in visual entrainment in neurotypical adults and provide novel evidence in favor of predictive coding accounts of autistic perception, as well as a consequent bias toward bottom-up sensory signaling over predictive feedback in a context of visual entrainment.

**Significance statement:** Traveling waves reflect the spatiotemporal propagation of neural oscillations, providing a window into hierarchical brain communication. By examining traveling wave dynamics during visual entrainment, we show that neurotypical adults exhibit increased backward (frontal-to-occipital) waves, consistent with enhanced top-down predictive signaling. In contrast, adults with Autism Spectrum Disorder (ASD) display a marked increase in forward (occipital-to-frontal) waves, indicating stronger bottom-up sensory drive. These findings provide electrophysiological evidence for atypical predictive processing in ASD, supporting predictive coding theories that propose an imbalance between sensory evidence and prior expectations. Our results highlight traveling waves as a sensitive neural marker of hierarchical signaling and predictive dynamics across typical and atypical perceptual systems.

## Introduction

Decades of research demonstrate the role of neural oscillations in various cognitive functions, including perception and sensory processes (Hipp et al., 2011; Arnal and Giraud, 2012; Samaha and Postle, 2015; VanRullen, 2016). Recent work has shown that oscillatory dynamics are best characterized considering their spatial component, as they propagate as traveling waves (Muller et al., 2018; Zhang et al., 2018). Importantly, oscillatory traveling waves have been related to several cognitive functions, including perception (Muller et al., 2014; Pang (庞兆阳) et al., 2020; Fakche et al., 2022), visual attention (Alamia et al., 2023), and episodic and working memory (Mohan et al., 2024; Zeng et al., 2024).

Considering scalp EEG recordings, several studies identified waves mainly traveling along the anterior-to-posterior axis (Alexander et al., 2013, 2019; Pang (庞 兆 阳) et al., 2020; Grabot et al., 2025): waves propagating from occipital to frontal regions (FW, forward waves) are more prominent during visual stimulation (Lozano-Soldevilla and VanRullen, 2019; Pang (庞 兆 阳) et al., 2020), whereas waves propagating in the opposite direction (BW, backward waves) dominate at rest, as well as during attentional suppression of visual input (Ito et al., 2005; Alamia et al., 2023). Interestingly, one study reported that during visual entrainment (i.e., when participants attended to rhythmic stimuli), the steady-state visual evoked potential (SSVEP) propagated as a wave from frontal to occipital regions in some subjects (Burkitt et al., 2000). Given the temporal predictability of rhythmic stimuli, these findings are consistent with the hypothesis that backward waves reflect top-down expectations (Samaha et al., 2015; Haegens and Zion Golumbic, 2018), as proposed within the predictive coding framework (Alamia and VanRullen, 2019; Friston, 2019; Schwenk and Alamia, 2025).

Traveling waves may provide insights into predictive processes, which are hypothesized to be atypical in several conditions. In particular, an imbalance in the weighting of predictions and prediction errors may underlie symptoms in Autism Spectrum Disorders (ASD) (Brock, 2012; Pellicano and Burr, 2012; Sinha et al., 2014; Van de Cruys et al., 2014). ASD is a neurodevelopmental condition characterized by deficits in social interaction and communication, repetitive behaviors, restricted interests, and atypical sensory responsivity (American Psychiatric Association, 2013). Predictive coding hypotheses suggest that perception in ASD may be less influenced by priors, either because priors are assigned low precision (Pellicano and Burr, 2012) or because sensory inputs are encoded with high precision (Brock, 2012). Alternatively, prediction errors may be particularly strong and inflexible in ASD (Van de Cruys et al., 2014), making the environment feel unpredictable and leading to sensory overload. However, empirical findings testing these hypotheses remain mixed, as prior precision in ASD has been reported as intact, increased, or decreased depending on the context (Cannon et al., 2021; Angeletos Chrysaitis and Seriès, 2023). In ASD, neural responses to prediction errors also appear more inflexible (Goris et al., 2018) and stronger in some brain regions (Sapey-Triomphe et al., 2023b).

In this study, we measure traveling waves to investigate sensory and predictive mechanisms further in both neurotypical and ASD populations. First, we aimed to replicate previous results (Burkitt et al., 2000), confirming the predominance of backward waves in a neurotypical population during visual entrainment. Then, considering TWs as markers of predictive coding processes (Alamia and VanRullen, 2019; Friston, 2019; Schwenk and Alamia, 2025), we hypothesize stronger forward waves in ASD than in neurotypical populations, consistent with predictive coding accounts favoring sensory inputs over priors. To test these hypotheses, we reanalyzed a previously published EEG dataset in which neurotypical and autistic adults perform a detection task during visual entrainment (Sapey-Triomphe et al., 2023a). Our results replicated previous findings in neurotypical individuals, revealing the predominance of backward waves during visual entrainment. In line with predictive coding accounts, we found the opposite pattern in autistic adults, with a substantial increase in forward waves in response to rhythmic stimulation compared with neurotypical adults.

## Material and Methods

### Participants

Data were collected from 25 neurotypical and 24 autistic adults between 18 and 54 years old, with normal or corrected-to-normal vision. Neurotypical participants were free of psychiatric or neurological disorders and scored below the cut-off of 32 on the Autism-spectrum Quotient (Baron-Cohen et al., 2001). Autistic participants were diagnosed based on the DSM-5 criteria (*Diagnostic and Statistical Manual of Mental Disorders (5th ed*.), 2013), at the Expertise Center for Autism in Leuven. Six autistic participants reported one or more comorbidities (e.g., attention disorders, dyslexia, dyscalculia, or depression). Due to a technical issue in the EEG recordings (i.e., an artifact at 16 Hz), we excluded two participants from the neurotypical group and six from the autistic group, resulting in 23 participants in the neurotypical group and 18 in the autistic group for the analyses. These groups were matched for age (NT: 30.8 ± 7.5; ASD: 30.1 ± 8.0), sex ratio (NT: 9 female; ASD: 8 female), handedness (NT: 3 left-handed; ASD: 3 left-handed), and total Intelligence Quotient based on the Wechsler Adult Intelligence Scale IV (2008) (NT: 110.3 ± 16.3; ASD: 109.4 ± 14.1).

According to the Helsinki Declaration, all participants provided informed, written consent before the study. This study was approved by the medical Research Ethical Committee of the university hospital KU/UZ Leuven.

### Experimental design

The dataset we analyzed is part of a larger study in which participants performed various tasks. We report the relevant information for this work below; for further details, please refer to Sapey-Triomphe et al. (2023a).

In this study, participants performed two tasks, each consisting of 24 trials. They were placed 90 cm from an LCD screen where the stimulus was displayed, in a darkened and quiet room. In both tasks, participants viewed a vertical sine-wave pattern-reversing grating with blurred edges (see Figure 1A), spanning 20° of visual angle, and presenting 20 pattern reversals per second (i.e., flipping between white and black lines). Overall, there were four types of trials: in half of the trials, the spatial frequency (SF) was set to 1.5 cycles per degree while the contrast increased or decreased following a logarithmic scale; in the other half of the trials, the contrast was fixed to 30% while the SF increased or decreased logarithmically. Each trial lasted 20 seconds, followed by 5 seconds of white noise stimulation covering the stimuli.

**Figure 1:**
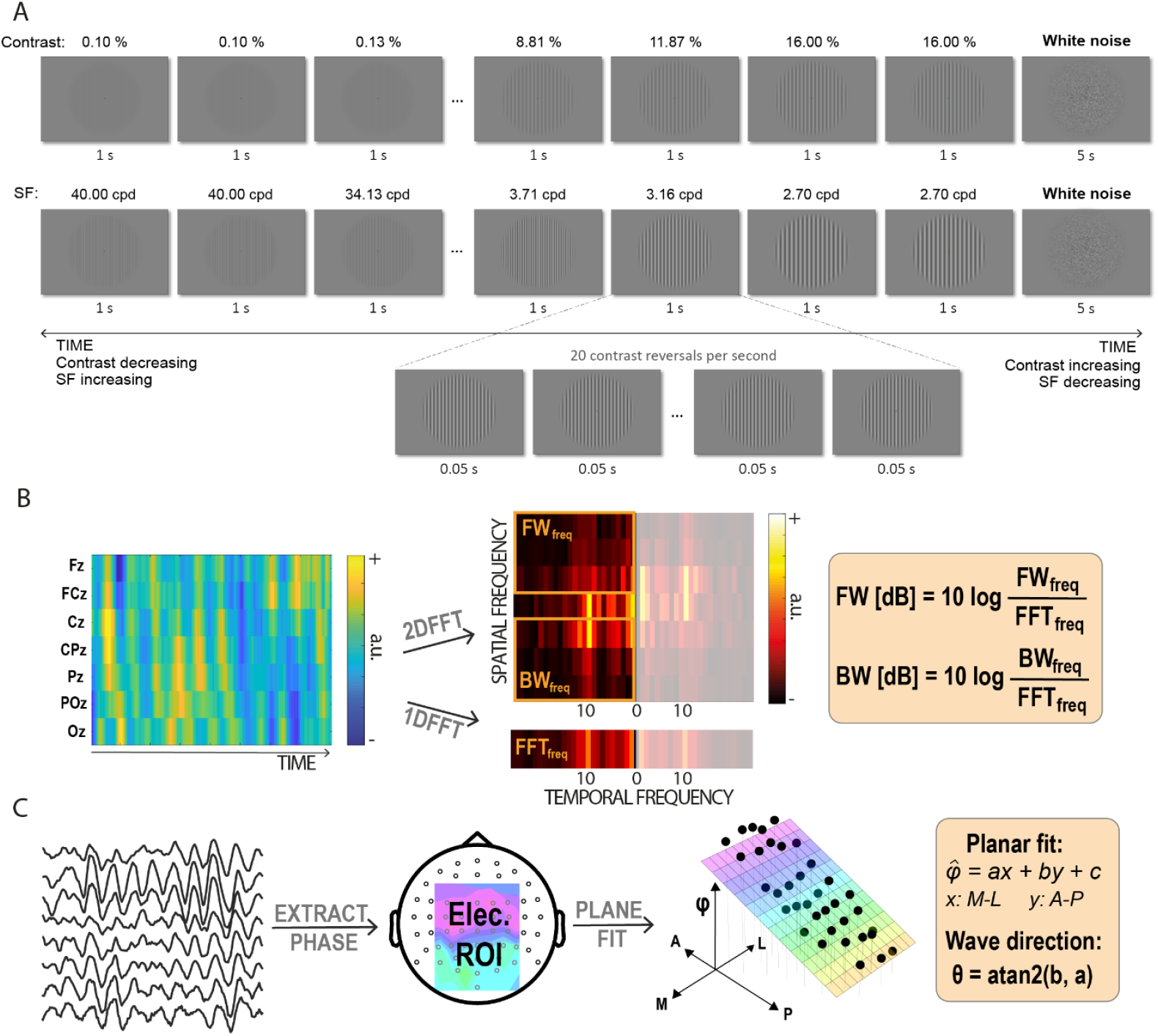
Experimental paradigm and wave computation methods. **A**. Experimental trials in which contrast-reversing gratings progressively increased or decreased in contrast (upper row) or spatial frequency (SF, lower row). The 20-second trials were separated by 5 seconds of white noise. The fast-periodic visual stimulation elicited a steady-state response at 20 Hz in the visual cortex (see Sapey-Triomphe et al., 2023a for further details). **B**. EEG data from seven midline electrodes is stacked into a 2D map, and transformed via a two-dimensional fast Fourier transform (2D-FFT). In the resulting power spectra, forward waves corresponded to the upper-left or lower-right quadrants, and backward waves to the lower-left or upper-right quadrants. For each temporal frequency and propagation direction, wave power was defined as the maximum value within the corresponding quadrant column, normalized by the mean FFT power at that frequency. **C**. For the alpha-band and the entrained frequency (i.e., 20Hz), the continuous phase is extracted from the EEG. Phase values across electrodes in the ROI are mapped topographically for each time point, and a plane is fitted to estimate parameters *a* and *b*, indicating wave direction.

In the first task, participants were asked to fixate on the central dot, while in the second task, they were asked to report a change in stimulus detection (perceiving the grating or not). Participants performed six trials per condition for both tasks, totaling 24 trials per task. The order of the trials was randomized during the experiment.

### EEG preprocessing

EEG data were recorded at 512 Hz using a 64-channel BioSemi Active 2 system with Ag-AgCl active electrodes. Electrode offset was kept below 30 mV. All preprocessing analyses were performed offline using custom MATLAB functions supported by the EEGLAB library (Delorme and Makeig, 2004). Specifically, we first downsampled the data to 160 Hz and then applied a notch filter, from 47 to 53 Hz, and a high-pass filter with a cut-off frequency of 1 Hz. We then re-referenced the data to the average of all electrodes. Lastly, we segmented the data into epochs starting 1 second before the onset of the stimulus and ending 1 second after the offset of the white noise.

### Traveling waves analysis

We performed two complementary analyses to quantify traveling waves during the task. The first method is based on the 2D Fast Fourier Transformation and has been used in previous studies investigating traveling waves in EEG signals (Alamia et al., 2024; Wei et al., 2024; Zeng et al., 2024). The second method, based on a linear fitting of the phases, was originally designed to detect waves in intracortical recordings (Zhang et al., 2018; Das et al., 2022) but has recently been adapted for surface EEG recordings (Schwenk and Alamia, 2025).

#### 2D-FFT analysis

For each trial, we extracted 1-second segments from the midline electrodes (i.e., Oz, POz, Pz, CPz, Cz, FCz, Fz) with a sliding window using a 500-ms overlap. These signals were arranged in 2D maps, with time and electrodes as axes. We applied a 2D-FFT to these images; the upper and lower quadrants represent the power of forward (FW) and backward (BW) waves as a function of temporal frequency. We extracted the values by averaging over the alpha band (8-12 Hz) and the entrained frequency (between 19 and 21 Hz). We then computed the amount of waves in decibels by considering the log ratio between these values and the averaged temporal power obtained using 1D-FFT transformation on each electrode separately.

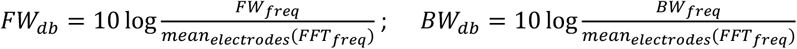

#### Phase fitting analysis

We first computed the continuous signal phase in the electrodes within a given region of interest (ROI), which included frontal, central, and parietal electrodes, separately in the alpha band (7-13 Hz) and between 19 and 21 Hz. Phases were estimated using the Hilbert transformation of the signal, which was bandpassed using an FIR filter. The phases were then referenced to the averaged phase within the ROI, and projected from the scalp to a 2D plane using the 10-20 layout provided by the FieldTrip toolbox (Oostenveld et al., 2011). We then estimated the phases using a linear plane fit given by:

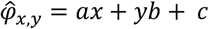

Where x, y are the electrodes’ positions, a, b the plane slopes, and c is the constant phase offset (average phase). For each time point, we evaluated the fit for a set of parameters defined over all possible directions θ, where θ is defined as θ = atan2(b, a). The best fit was selected by maximizing the vector length of the summed residuals in the circular space. Additionally, for each fit, we computed the proportion of variance explained by the fit (*ρ*^2^_cc_) from the circular correlation between predicted and observed phases as a measure of the goodness of fit. We then computed a null distribution of *ρ*^2^_cc_ values in a bootstrap fashion by randomizing the electrodes’ position 10 times (obtaining a single null distribution for each subject). For the actual data, we then considered only fits that fell above the 99^th^ percentile of the null distribution. Over conditions and at each time point, we computed the proportion (across trials) of FW and BW waves (defined by *α* equal to 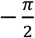 and 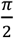 respectively, with a tolerance of 0.5 radians).

#### Granger analysis

We performed connectivity analysis using Granger causality in the spectral domain, computed using the MVGC toolbox (Barnett and Seth, 2014). For each condition, we considered the 4 seconds with the largest response amplitude (at the beginning or end of the trial, depending on the condition, when the grating is the most visible) to estimate Granger Causality (GC) values. We estimated the order of the model for each participant separately, optimizing the Akaike Information Criterion (with a maximum order of 20) and setting the maximum autocovariance lag to 3000. We ran this analysis over the midline electrodes (Oz to Fz). To obtain final estimates for FW- and BW-directed information flow, we averaged all GC values for pairs in one direction (e.g., for FW: Oz-POz, Oz-Pz, POz-Pz, etc.).

### Statistical analysis

All analyses were computed using Bayesian repeated-measure ANOVA, thus providing Bayes factors for each considered variable. Bayes Factors (BF) quantify the ratio between the models in favor of the alternative against the null hypothesis. In this work, we report all BFs having the alternative hypothesis at the numerator (conventionally BF_10_). We considered the amount of traveling waves in dB or the wave’s probability as the dependent variable, as well as different dependent factors and their interactions. The GROUP factor, which distinguishes between the neurotypical and autistic populations, was considered a between-subjects factor across all analyses. All analyses were implemented in JASP (Love et al., 2019), considering default uniform prior distributions.

## Results

In this study, we investigated how oscillations propagate through the cortex during a visual entrainment task in both neurotypical and autistic groups. Considering this novel perspective based on oscillatory traveling waves, we reanalyzed a previously published EEG dataset (Sapey-Triomphe et al., 2023a) in which participants viewed a grating with a 20 Hz pattern reversal while either the contrast or spatial frequency increased or decreased over 20 seconds. We quantified traveling waves using two complementary methods (Figure 1B and C), focusing on alpha-band oscillations and the entrained frequency (i.e., 20Hz).

### Traveling waves during visual entrainment and white noise

First, we quantified the amount of traveling waves propagating from the occipital to the frontal regions (forward) or in the opposite direction (backward, from the frontal to the occipital areas) using the 2D Fourier transform (2D-FFT) method (Alamia et al., 2024; Zeng et al., 2024). This approach provides power spectra of waves propagating along a given line of electrodes in both directions (in our study, we chose the electrodes along the midline, from Oz to Fz – see Figure 1B). We quantify traveling waves across the entire spectrum (Figure 2A), but focus our analysis on the entrained frequency (20 Hz) and the alpha band (8-12Hz, Figure 2B), which are supposedly reflective of predictive coding processes (Han and VanRullen, 2017; Alamia et al., 2024; Gabhart et al., 2025). The left panels of Figure 2A show the results for forward and backward waves in both groups during the 20 Hz visual entrainment (i.e., during the 4 seconds when the grating was the most visible): we found a clear peak in the spectrum of the backward waves (propagating from frontal to occipital regions, lower panel) at the stimulation frequency (i.e., 20 Hz) in the neurotypical (NT) group. Even though previous studies showed that forward waves have been associated with visual stimulation in EEG recordings (Pang (庞兆阳) et al., 2020; Alamia et al., 2023), our results are in line with a previous study showing a similar phase pattern during visual entrainment (Burkitt et al., 2000) Notably, the backward peak at 20 Hz was not present during the five-second visual stimulation with white noise (right panels of Figure 2A), and was not present in the spectrum of the waves propagating in the forward direction. Quite remarkably, however, we found a different pattern of results in the autistic (ASD) group. Specifically, we reported a smaller peak in the backward waves compared to the NT group, and a larger peak in the forward waves. As expected, we did not observe a response at 20 Hz during the presentation of the white noise stimulus. Overall, our results were supported by a repeated-measures Bayesian ANOVA, in which we tested the effects of STIMULI (entrainment vs. white noise), DIRECTION (forward vs. backward), and GROUP as between factors (ASD vs. NT groups), and are represented in Figure 2B. All factors revealed a large BF (STIMULI and DIRECTION: BF_10_ > 10^3^, GROUP: BF_10_ = 37), as well as their interactions (STIMULI x DIRECTION and STIMULI x GROUP: BF_10_ > 17, DIRECTION x GROUP: BF_10_ = 162; triple interaction STIMULI x DIRECTION x GROUP: BF_10_ = 108). Post-hoc analyses confirmed a strong difference between GROUPS during visual entrainment (BF_10_=72), but not during white noise (BF_10_ = 0.4), and similar results for the interaction DIRECTION x GROUP during entrainment (BF_10_ > 300) but not during white noise (BF_10_ = 1).

**Figure 2:**
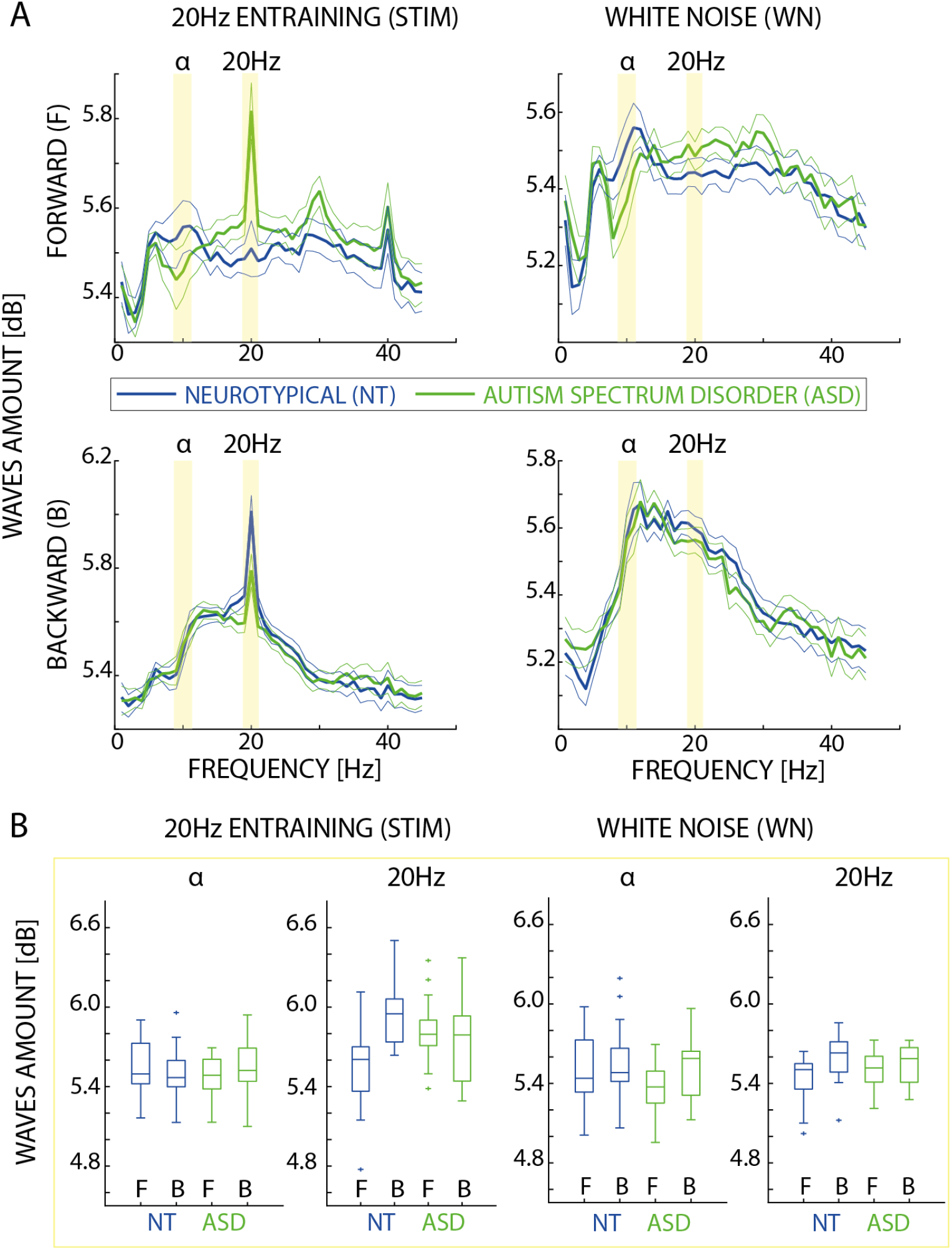
Full spectra traveling waves analysis based on 2D-FFT method. A. Spectra of forward and backward traveling waves in the upper and lower panels, respectively. The left column shows the traveling waves during the four seconds of highest visual entrainment, while the right column shows the waves during the five seconds of white noise. The averages and standard errors of the neurotypical (NT) group are represented in blue, while those of the ASD group are represented in green. The yellow bands indicate the interval used to extract the values in the alpha band and around the entraining frequency. B. Boxplots for the two groups during visual entrainment (left panels) and during the white noise (right panels), for both the alpha-band and the entraining frequencies.

Concerning the waves in the alpha band, we tested the difference between the NT and ASD groups during entrainment and white noise stimulation. A repeated-measure Bayesian ANOVA, using the same factors as for the 20 Hz frequency band, did not reveal any noticeable effect (all factors and interactions: 0.4 < BF_10_ < 1.1, except for STIMULI x DIRECTION, BF_10_ = 3.0).

### Differences between conditions and time courses

The visual stimulation task consisted of four types of trials: either the Gabor patch contrast or its spatial frequency was increased or decreased over a period of twenty seconds. We then analyzed how the amount of traveling waves changes as a function of the increase (or decrease) in the entraining visual stimulation, as shown in Figure 3A. Regarding the backward waves, we observed a larger modulation over time in both groups, particularly in conditions where the spatial frequency changed over time. A repeated-measure Bayesian ANOVA confirmed such results, providing very large BFs for the factors CONDITION (for the four types of trials), TIME (composed of 5 BINS of 4 seconds each), and their interactions (all BF_10_ > 10^13^). However, we did not observe any relevant difference between GROUP (BF_10_ = 1.5) or any interactions (BF_10_ < 0.02). On the other hand, we observed a substantial difference between GROUPs when considering the forward waves, as well as the factors CONDITION, TIME, and all the interactions (all BF_10_ >10^10^ for the factors, and BF_10_ >10^5^ for the interactions). These results suggest that the temporal dynamics of visual entrainment elicit larger forward waves in the ASD group, specifically in spatial frequency conditions.

**Figure 3:**
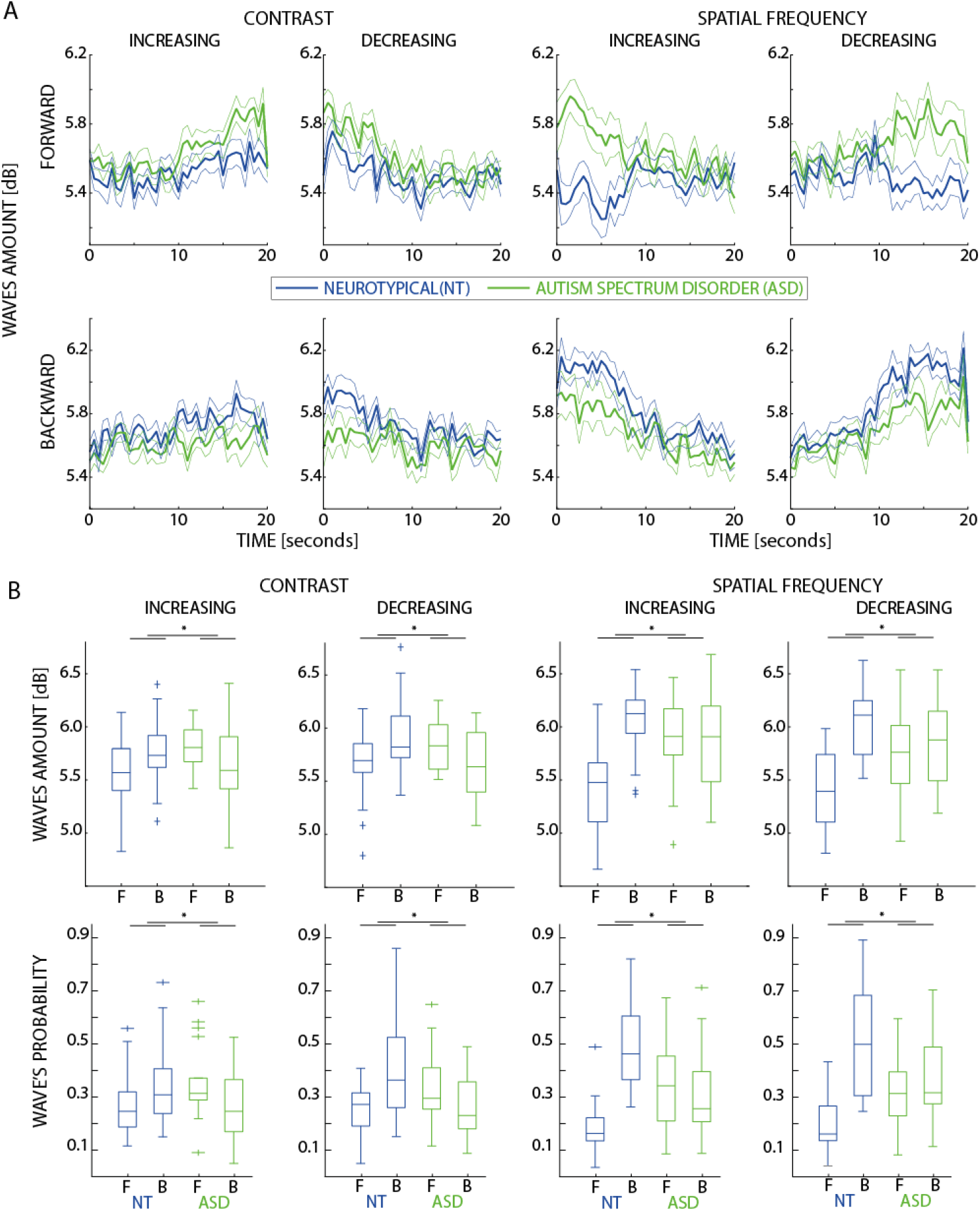
Traveling waves entrained at 20 Hz over time (2D-FFT method). A. Traveling waves for both neurotypical (NT, in blue) and ASD (in green) groups in all four conditions (columns) for forward and backward waves (upper and lower panels, respectively). In each condition, either the contrast or the spatial frequency increased or decreased over twenty seconds. B. Boxplots with the average and standard errors of traveling waves for the four seconds with the highest entrainment (either the first or the last four seconds). The first row shows the waves computed using the 2D-FFT method, while the second row shows the waves calculated using the phase plane fitting method. Asterisks highlight the effect in the GROUP x DIRECTION interaction.

In addition, we analyzed the differences between conditions for both forward and backward waves at 20 Hz for both groups in the 4 seconds with the highest entrainment (either at the beginning or at the end of the trial, depending on the condition, Figure 3B). A repeated-measure Bayesian ANOVA revealed a substantial effect for the factors CONDITION (BF_10_ > 5*10^3^), DIRECTION (BF_10_ > 2.9*10^5^), as well as GROUP (BF_10_ = 6.9). We also found a significant interaction for the CONDITION x DIRECTION (BF_10_ > 2.5*10^4^) and GROUP x DIRECTION (BF_10_ = 29.8), whereas all other interactions had 0.4 < BF_10_ < 0.8.

### Results of the phase fitting analysis

To further corroborate our results, we replicate the same statistical analysis using a different method to quantify traveling waves. Specifically, we estimated the direction of traveling waves by fitting a plane to the estimated phase of a given region of interest, centered around the midline (see Methods for details). Unlike the 2D-FFT method, which may incorporate residual amplitude modulations into its estimate despite its baseline, the phase fitting method relies solely on the analytic phase of the signal, thereby eliminating potential confounding effects due to oscillatory power. The Bayesian ANOVA performed on the direction calculated from the phase fit confirmed all previous results, specifically a GROUP × DIRECTION interaction (BF_10_ = 15.6), but not a triple interaction (GROUP × DIRECTION × CONDITION: BF_10_ = 0.29). Altogether, these results confirm a difference between the ASD and NT groups across conditions, with an increase in forward waves in the ASD group induced by the visual entraining stimulation.

### Forward and backward wave propagation stable-state duration

A critical advantage of the phase-fitting method, compared to the 2D-FFT method, is the better temporal resolution, as it estimates the wave’s direction at every time point from the analytical phase. Taking advantage of this improved temporal resolution, we performed an additional analysis to assess the duration of continuous time periods (‘states’) during which we reliably observed forward or backward waves at 20 Hz during the visual entrainment task. Figure 4 illustrates the difference between groups for both directions. A repeated-measure Bayesian ANOVA revealed a substantial difference between DIRECTIONS (BF_10_ = 103.1), between GROUPS (BF_10_ = 16.5), and their interactions (BF_10_ = 58.4). These results indicate that forward wave states lasted longer in the ASD group than in the NT group, and similarly, albeit to a lesser extent, for backward waves. We then plot the distribution of state durations for both groups and directions, and fit an exponential function for each subject as:

**Figure 4:**
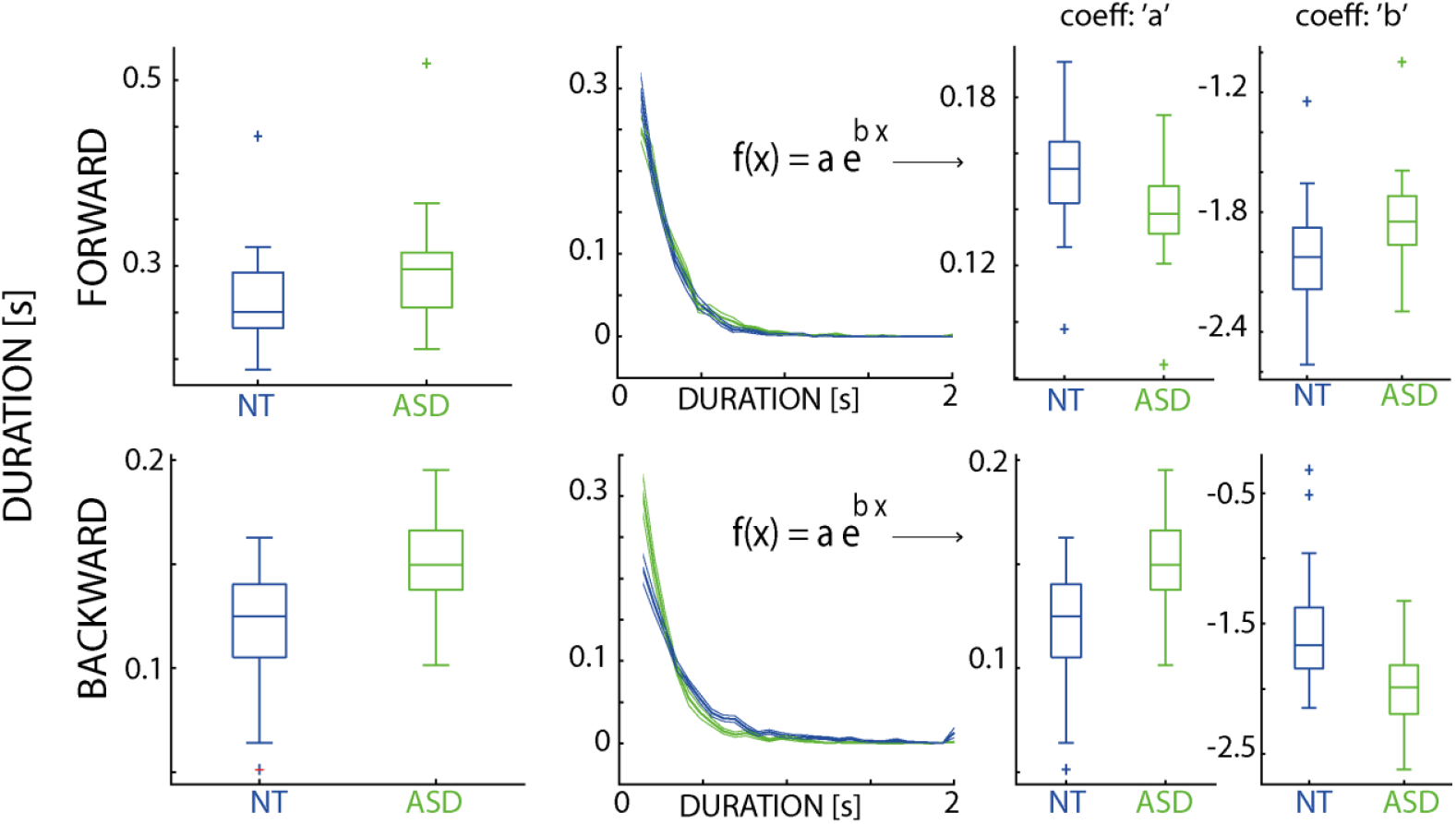
Traveling waves state analysis, based on the plane fitting method. The first column shows the boxplots for the temporal duration of forward and backward traveling waves, estimated via the phase fitting method. The second column illustrates the exponential fitting performed over the duration for the two groups: neurotypical (NT) in blue and ASD in green. The last columns show the boxplots of the fitted coefficients: the amplitude ‘*a*’ and the time constant ‘*b*’.

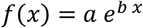

where the ‘*a*’ and ‘*b*’ coefficients characterize the amplitude and time constant of the function, respectively. As shown in Figure 4 and corroborated by two repeated-measure Bayesian ANOVAs, for both coefficients, we observed a substantial difference between the two GROUPs (‘*a*’: BF_10_ = 53.4; ‘*b*’: BF_10_ = 45.9) and for the two DIRECTIONs (‘*a*’: BF_10_ = 221.8; ‘*b*’: BF_10_ = 199.7). We also reported for both coefficients a significant interaction GROUP x DIRECTION (‘*a*’: BF_10_ = 216.1; ‘*b*’: BF_10_ = 179.9), suggesting shorter time constants for the ASD group (confirming an overall longer duration), and smaller intercept in the forward direction, and the opposite pattern in the backward case.

### Granger analysis

Lastly, we investigated whether the increase in backward and forward traveling waves in the neurotypical and ASD groups, respectively, was also observed using different causal measures of connectivity. In particular, we performed frequency-resolved Granger Causality (GC) connectivity during the visual entrainment task, which revealed a difference between the two groups at the entrained frequency band (around 20 Hz, 19-21 Hz) and in the alpha band (8-12 Hz). Previous theoretical work, based on linear autoregressive models, demonstrated that phase-based measures may reveal an opposite direction than the one observed with causal methods (Alamia et al., 2025), particularly when the connections in the same direction of the wave are inhibitory. Figure 5 shows the mean results for both groups, averaging the pair of consecutive electrodes along the midline in the occipital-to-frontal direction (forward) and vice versa (backward). Unlike the traveling waves analysis, the GC results did not corroborate any difference between the two GROUPS (BF_10_ = 1.5) or any interactions between any factor and GROUP (all 0.9 < BF_10_ < 1.1). Still, they showed a difference between DIRECTIONS (BF_10_ = 22.5) and BANDS (BF_10_ = 19.9), as well as their interaction (BF_10_ = 94.8). All in all, these results suggest that the traveling wave and the Granger analysis provide different results as only in the former case we observed a difference between groups.

**Figure 5:**
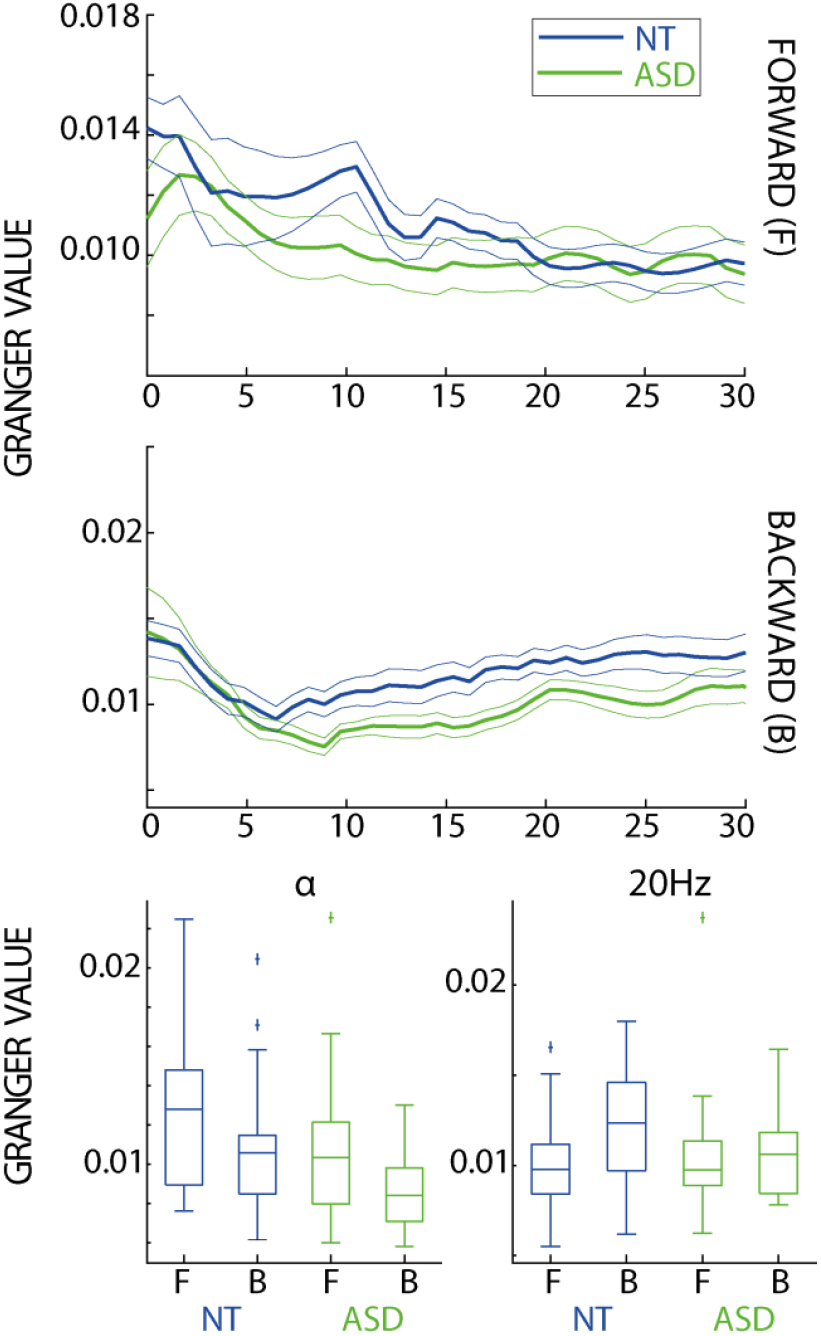
Granger analysis over midline electrodes. Results of the spectral Granger analysis for the neurotypical (NT) and ASD groups in the occipital to frontal (forward) and opposite (backward) directions. The boxplot displays the results for the alpha band and the entrained frequency.

## Discussion

In this study, we quantified oscillatory traveling waves to investigate sensory and predictive mechanisms in both neurotypical and autistic (ASD) populations. Consistent with previous findings (Burkitt et al., 2000), we observed enhanced backward waves in neurotypical individuals during visual entrainment, at the frequency of the SSVEP. In contrast, individuals with ASD demonstrated stronger forward waves relative to neurotypical participants, aligning with predictive coding accounts of ASD that posit an imbalance favoring sensory inputs and prediction errors over prior expectations. Our findings extend previous research linking occipital-to-frontal wave propagation to visual perception and sensory processing (Alamia and VanRullen, 2019; Lozano-Soldevilla and VanRullen, 2019; Pang (庞兆阳) et al., 2020; Wei et al., 2024), as well as studies associating traveling waves with predictive processes (Alamia and VanRullen, 2019; Friston, 2019; Alamia et al., 2024).

In this work, we consider neural visual entrainment as the synchronization (or phase-locking) of neural oscillations with external rhythmic stimuli (Lakatos et al., 2019). This phenomenon has been extensively studied using scalp-recording techniques with visual stimuli (Mathewson et al., 2012; Cravo et al., 2013; Graaf et al., 2013), and previous work has shown that early visual cortical areas play a key role in generating the entrained signal (Spaak et al., 2014). However, other authors have proposed that neural entrainment represents a general mechanism shared across brain regions, enabling the coupling between internal and external (both exogenous and interoceptive) stimuli (Lakatos et al., 2019). In this view, entrainment serves as a means of broadcasting information within the brain. Consequently, neural entrainment may function as a dynamic process for selecting and coordinating neuronal excitability across brain regions (Helfrich et al., 2019; Obleser and Kayser, 2019; Duecker et al., 2024), as well as for enhancing the processing of sensory information that occurs at specific phases within a temporal structure (Jones et al., 2002; Nobre et al., 2007). In this sense, sensory entrainment can be naturally interpreted within the predictive coding framework, as it supports temporal prediction, allowing the brain to anticipate upcoming stimuli by aligning neuronal excitability with the expected onset of stimuli (Beker et al., 2021; Cannon, 2021; Doelling et al., 2023). In predictive coding terms, oscillatory alignment minimizes prediction error by timing cortical responses to expected events.

Interestingly, different frequency bands have been related to distinct predictive coding processes. Regarding traveling waves, alpha-band oscillations have been associated with both predictions and prediction errors (Alamia and VanRullen, 2019; Alamia et al., 2024; Schwenk and Alamia, 2025). More generally, slower frequencies have been associated with top-down processes, whereas gamma oscillations have been linked to prediction errors (Michalareas et al., 2016; van Pelt et al., 2016; Strube et al., 2021; but see Vinck et al., 2025). Regarding neural entrainment, different cortical areas exhibit frequency-specific resonance properties, for example, in the alpha band, which have been shown to have the strongest neural entrainment compared to other frequency bands (Herrmann, 2001). Future work may explore how entrainment influences perception and propagates through brain regions at slower frequencies (i.e., alpha or theta bands), thereby targeting, for example, specific top-down mechanisms.

In this study, participants actively attended the flickering visual stimuli to perform a discrimination task (Sapey-Triomphe et al., 2023a). Previous studies have shown that neural entrainment, although primarily an automatic process, can be modulated by top-down mechanisms, which enhance or suppress entrainment depending on task demands (Mathewson et al., 2012; Calderone et al., 2014; Gray et al., 2015; Helfrich et al., 2019). On the one hand, one might expect similar but weaker results during passive, task-free perception of the same stimulus. On the other hand, we did not find any between-subject correlation between forward or backward waves and different measures of task accuracy (i.e., contrast and spatial frequency visual thresholds; data not shown, all p-values>0.05), suggesting that oscillatory traveling waves may not reflect behavioral performance.

Regarding the ASD group, our results revealed an atypical pattern of oscillatory traveling waves, which is hypothesized to reflect differences in predictive processing. Specifically, autistic participants showed increased forward waves relative to backward waves, whereas the NT group displayed the opposite pattern. Within the predictive coding framework, this suggests a more substantial weighting of sensory inputs and prediction errors in autistic adults in this context of visual entrainment, consistent with theories positing a reduced influence of priors relative to sensory evidence in ASD (Brock, 2012; Pellicano and Burr, 2012). This finding also aligns with the HIPPEA account, which hypothesizes High and Inflexible Precisions of Prediction Errors in ASD (Van de Cruys et al., 2014). Such mechanisms might also contribute to the stronger neural responses to mid-level precision-weighted prediction errors observed in a few brain regions in ASD (Sapey-Triomphe et al., 2023b).

Group comparisons further revealed increased forward and reduced backward waves in autistic participants during visual entrainment as compared to NT participants. Interpreted through the lens of predictive coding, these results would reflect stronger prediction errors and weaker priors in ASD than in NT participants. The group differences were most pronounced for trials varying in spatial frequency, and to a lesser extent for those varying in contrast. Interestingly, psychophysics tasks in the same sample showed that autistic participants exhibited higher detection and responsivity thresholds for spatial frequency than NT, while no significant group differences emerged for contrast (Sapey-Triomphe et al., 2023a). Because higher spatial frequencies are intrinsically harder to detect, these behavioral results suggest sensory hypersensitivity and overresponsivity to spatial frequency in ASD. The observed increase in forward waves may thus reflect stronger propagation of sensory inputs and prediction errors, contributing to this visual hypersensitivity and overresponsivity in ASD. Note that a tendency to rely more heavily on high spatial frequencies may bias perception toward local details in ASD, and thereby contribute to specificities in face processing (Deruelle et al., 2004; Kéïta et al., 2014). Overall, our findings on traveling waves indicating a stronger reliance on sensory evidence than on priors could contribute to the self-reported visual hypersensitivity frequently described in ASD (Simmons et al., 2009; Tavassoli et al., 2014; Robertson and Simmons, 2015; Dellapiazza et al., 2018; Sapey-Triomphe et al., 2023a).

This atypical balance in traveling waves in favour of forward waves can further be interpreted within the effective connectivity literature, which reported enhanced bottom-up and/or diminished top-down connectivity during perceptual and learning tasks in ASD, often associated with an increased activation in sensory areas (e.g., Sapey-Triomphe et al., 2020; Randeniya et al., 2023). It may also reflect or contribute to the decreased habituation to sensory inputs in ASD (Jamal et al., 2021; Dwyer et al., 2023).

Overall, our results provide new insights into the role of oscillatory dynamics in sensory processes, particularly in visual entrainment. The predictive coding interpretation offers a compelling theoretical framework for interpreting and modeling brain dynamics (Friston, 2019; Alamia et al., 2024). Future work will test this framework more directly, using, for example, experimental design based on statistical learning or expectation violation. Replicating and extending the results with complementary imaging measures, such as MEG recordings, could provide better spatial resolution and overcome some limitations that affect traveling wave analyses in EEG recordings, such as the disruption of long-range connections, signal smearing (Nunez, 1974; Alexander et al., 2019), and the interpretation of the source origins of traveling waves signals (Zhigalov and Jensen, 2023; Petras et al., 2025).

## Acknowledgments

The authors thank the participants for their time, Naomi Couder for her help in recruiting participants, and Joke Dierckx for her help in acquiring data. A.A. was funded by the European Union under the European Union’sHorizon 2020 research and innovation program (grant agreements No. 101075930). The copyright holderfor this is of the author(s) only and do not necessarily reflect those of the European Union or theEuropean Research Council (ERC). Neither the European Union nor the granting authority can be heldresponsible for them.

